# Cervical Mucus Viscoelasticity and Sperm Velocity are Correlated and Concentration-dependent *In Vitro*

**DOI:** 10.1101/2025.05.18.654742

**Authors:** Matthew R. Markovetz, Shuhao Wei, Chris Celluci, Mackenzie Roberts, Leo Han

**Affiliations:** Marsico Lung Institute, University of North Carolina at Chapel Hill, Chapel Hill, USA; Department of Obstetrics and Gynecology, Oregon Health Sciences University, Portland, USA; Oregon National Primate Research Center, Division of Reproductive and Developmental Sciences, Portland, USA

## Abstract

**Background:** Mucus in the endocervix serves as fertility gatekeeper in the reproductive tract through hormonally regulated changes in biophysical properties. Cervical mucus can thicken to prevent ascension of sperm into the upper reproductive tract or thin to permit fertilization. Current reproductive studies of mucus viscoelastic properties rely on subjective visual appraisal of cervical mucus changes. Our goal was to use particle tracking microrheology (PTMR) to objectively assess cervical mucus viscoelastic properties and associate these measurements with in vitro measures of sperm velocity.

**Methods:** Using cervical mucus obtained from rhesus macaques (*Macaca mulatta*) at necropsy, we used to PTMR to measure viscoelasticity (η*) under stepwise, serial dilutions. In parallel we measure sperm velocity using custom sperm tracking and analysis workflows.

**Results:** We report that both mucus η* and sperm velocity displayed a concentration-dependent behavior, where η* increased as mucus concentration increased, and sperm velocity correspondingly decreased. Viscoelasticity and sperm velocity were strongly negatively correlated (p<0.001).

**Conclusions:** PTMR and sperm tracking in mucus provide quantitative measure of viscoelastic mucus changes. PTMR is potentially a method for quantitatively assessing fertility potential in the cervix that could be applied to both infertility and contraceptives studies.

## Introduction

Cervical mucus is a critical regulator of female fertility^1^ and a natural immune barrier to the upper female genital tract.^2^ Mucus biophysical properties fluctuate during the menstrual cycle in response to changes in sex-steroid hormones; mucus can decrease in viscoelasticity during the peri-ovulatory time period to permit sperm ascension into the reproductive tract and increase during other time points to create a thick sperm-impenetrable barrier.^3^ In fact, hormonal contraception, in particular progestin-only contraception, relies on this mechanism to maintain high efficacy even when ovulation occurs.

While we understand many of the hormonal regulators and drivers of mucus changes, we have limited tools for measuring observed changes in cervical mucus. The current standard for clinical evaluation of cervical mucus in contraceptive and fertility studies is the cervical mucus score (CMS aka Insler score).^4^ Insler scoring is only semi-quantitative as it summation of 5 qualitative assessments of mucus characteristics (ferning, spinnbarkeit (stretchability), viscosity qualitative), cellularity, quantity). While widely used in clinical trials, insler score is limited in objectivity, reproducibility, and interpretability of the final score as the socring system has never been directly validated for fertility.^4^ Similarly, other measures of mucus that assess the sperm-mucus interaction lack standardized methodologies and are often difficult to interpret.^5^ As a result, there remains a need for a robust, quantitative assay of cervical mucus biophysical properties that is related directly to fertility and sperm function.

Prior studies have shown that cervical mucus viscoelasticity and penetrability by sperm are inversely related, with minimum viscosity and maximum penetration occurring during the ovulation period.^3,6^ Additional studies, primarily in pulmonary mucus, have shown that concentration is the primary determinant of mucus viscoelasticity,^7^ and several early works on cervical mucus rheology reported that while cervical mucus volume and production is highest during ovulation, its concentration is lowest mid-cycle.^6,8^ Our hypothesis in this work is that both cervical mucus viscoelasticity and sperm penetrability are mucus concentration-dependent phenomena. Using cervical mucus and sperm obtained from a non-human primate model, we demonstrate that viscoelasticity and sperm motility are inversely related when quantified using particle tracking methodologies.

## Materials and Methods

### Non-human primate cervical mucus and sperm collection

Cervical mucus and sperm were obtained from rhesus macaques (*Macaca mulatta*) housed at the Oregon National Primate Research Center (ONPRC) in Portland, Oregon. Mucus was obtained at the time of necropsy from adult female macaques as previously described.^9^ For sperm collection, all males were trained by the ONPRC Behavioral Services Unit for collaborative semen collection by non-sedated electro-ejaculations.^10^ Semen samples were collected and allowed to liquefy at 37°C for 30 min before evaluation. The liquid fraction of the sample was washed and resuspended with warm HEPES-buffered Tyrode albumin lactate pyruvate (TALP-Hepes) with bovine serum albumin (BSA) supplemented at 3 mg/ml resuspended in the remaining 1 ml. All work was carried out in strict accordance with the recommendations in the Guide for the Care and Use of Laboratory Animals of the National Institutes of Health. The ONPRC Animal Care and Use Committee reviewed and approved all animal procedures.

### Particle tracking microrheology and sperm penetration

Rheological measurements of cervical mucus were performed using particle tracking microrheology (PTMR).^11^ Cervical mucus samples were prepared at concentrations ranging from their harvested value and at several diluted values obtained via serial dilution of aliquots into equivalent volumes of Talp-Hepes. Carboxylated, 1 μm, polystyrene beads (Fluospheres, Thermo Fisher, Fremont, CA, USA) were added to mucus samples at a dilution of 1:600. Beaded samples were placed in a sample chamber prepared by cutting out a small square from a strip of Parafilm placed on a glass slide (Fisher). All samples were performed in technical duplicate to account for variability in each aliquot. The chambers were sealed by placing a cover slip over the open portion of the Parafilm. Thermal diffusion of the beads in mucus was recorded for 10 seconds at 30 frames per second using an Insight Gigabit camera (Spot Imaging) on an inverted light microscope (Zeiss AX10). Bead motion was tracked via a custom Python program using the TrackPy package (https://doi.org/10.5281/zenodo.34028). The mean squared displacement of bead diffusion was converted into viscoelastic moduli in accordance with previous methods.^12^

Semen was added to parallel aliquots of the same mucus samples prepared and diluted for PTMR at concentration of 10^6^ sperm/ml. Sperm were fluorescently labeled with Hoescht in order to improve trackability. Using identical sample chambers as those used in PTMR experiments, videos of sperm motion in mucus were recorded for 5 seconds at 20 frames per second. Sperm motion was tracked using a slightly modified version of the PTMR program to account for the larger size of sperm and their faster, directed motility. Straight-line velocity was calculated for all sperm trajectories with recorded duration of at least one second. All experiments were performed with technical triplicates.

### Statistics

Linear model assessment was performed using the Matlab (© 2022 The MathWorks, Natick, MA, USA) function *corr*, which automatically computed the correlation coefficient and p-value of the model. Anderson-Darling tests to determine the appropriate family of distributions for ensemble data in PTMR and sperm were performed using the Matlab function *adtest*.

## Results

### Cervical mucus rheology is concentration-dependent

Mucus samples from n=6 female macaques were obtained at necropsy at differing points in the menstrual cycle and aliquoted and diluted serially up to dilutions as high as 1:128. The complex viscosity (η*) of these samples was measured using PTMR with HEPES buffer (η* = 10^−3^ Pa·s) serving as a reference control. Figure 2A illustrates the general viscoelastic behavior of mucus and sperm motility in mucus as sample dilution increased. Despite some variation due to inherent sample heterogeneity, we identified a trend that decreasing concentration decreased viscosity of the cervical mucus, which is consistent with mucus from other tissues.^13^ Due to low sample volumes, absolute concentration in terms of % solids was not determinable. As such, viscosities are reported with respect to relative dilution from the initially obtained sample.

**Figure 1.**
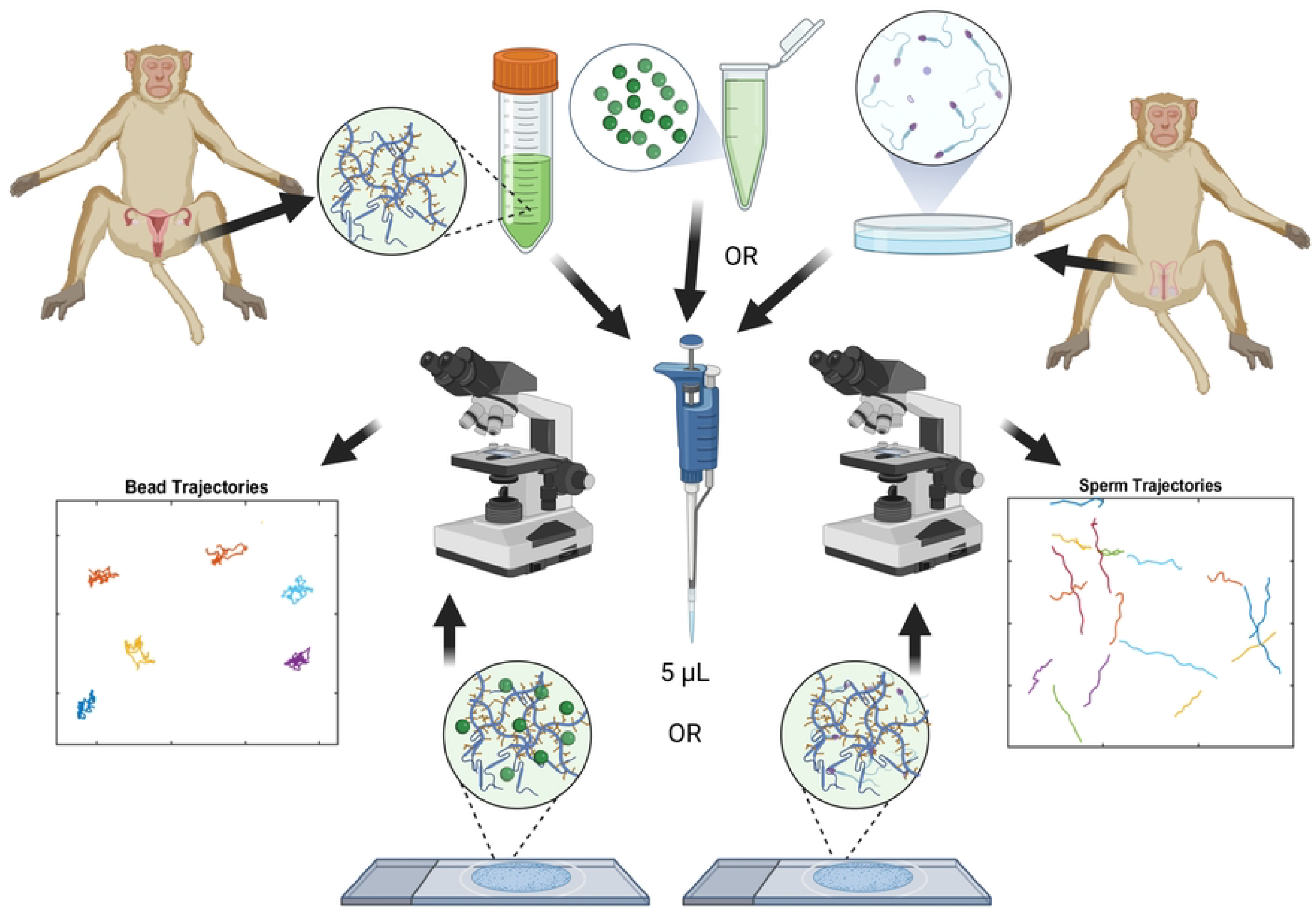
Schematic of our parallel assessment of sample microrheology and sperm motility in cervical mucus. Created partially in BioRender. Markovetz, M. (2025) https://BioRender.com/idydjtu

**Figure 2.**
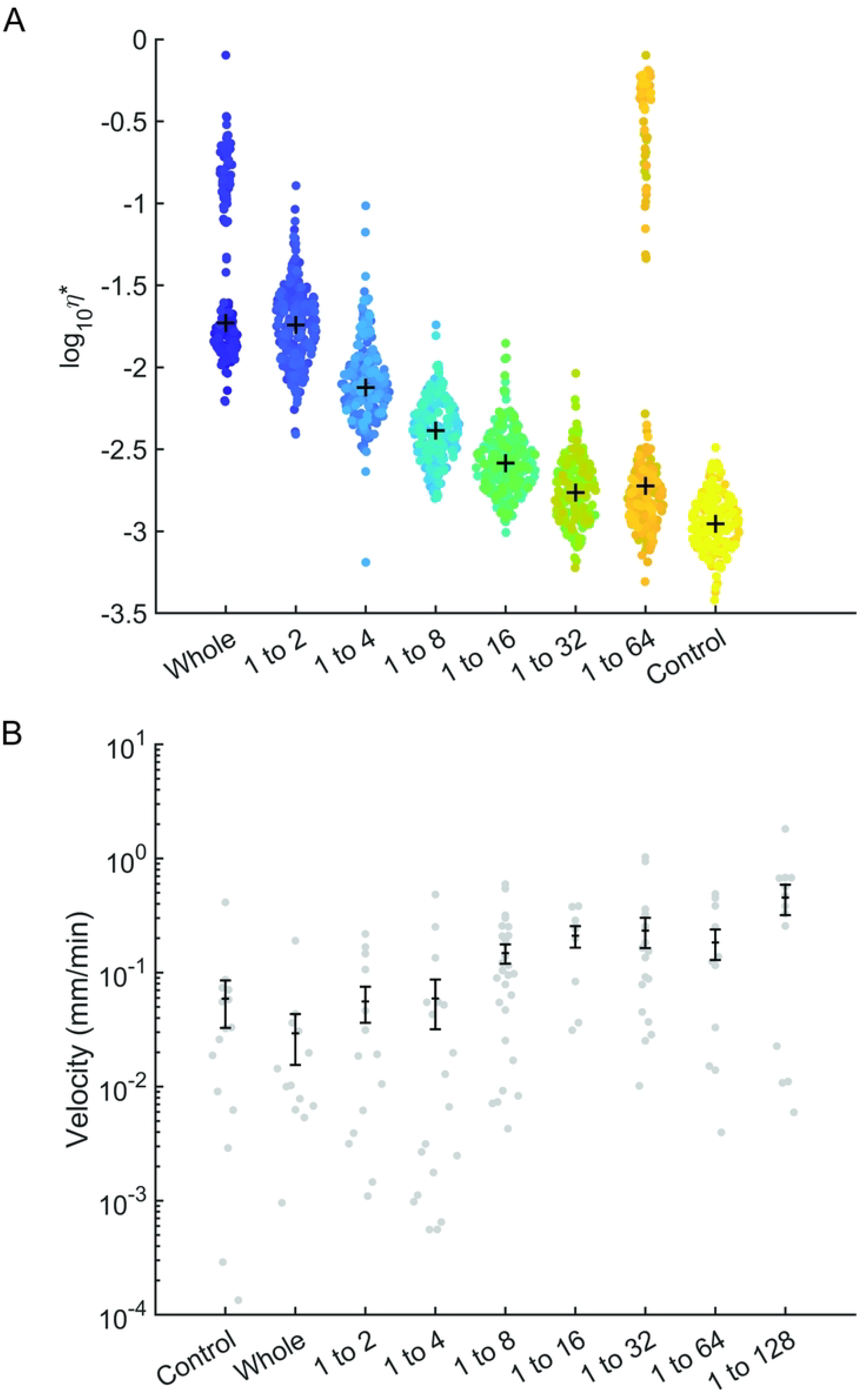
Representative ensemble behavior observed from (A) PTMR and (B) sperm velocity assessment across a range of serial dilutions. Mucus complex viscosity decreased, and sperm velocity increased, as mucus samples became more dilute.

### Sperm velocity within cervical mucus is concentration-dependent

Washed sperm was added to parallel aliquots of the mucus used in PTMR experiments. Sperm tracking in mucus was performed in identical experimental chambers as PTMR as most mucus specimens were too viscous to be loaded into a traditional computer assisted sperm analysis (CASA) device.^5^ Supplemental Video S1 gives an example video of sperm motion in cervical mucus with annotations for successfully tracked spermatozoa. Figure 2B shows the linear velocity of sperm in various concentrations of mucus from a single animal. This sample was representative of overall behavior seen in all animals, where velocity appeared to increase as mucus concentration decreased. Ultimately, decreasing cervical mucus concentration increased sperm velocity scaling with a power law relationship of ∼ c^0.5^. Velocity was highly variable within each sample, illustrating the heterogeneity of both mucus biophysics and sperm motility. Changes in response to dilution were not observed in other CASA parameters such as straightness, linearity, and average lateral head displacement. Sperm velocity in the most dilute samples exceeded that of sperm in buffer control, which reflects the known phenomenon that cellular swimmers can be faster in viscoelastic media compared to purely viscous media.^14^ *Sperm velocity is correlated with mucus viscoelasticity*

Because PTMR and sperm motility measurements were performed in parallel, the relationship between mucus rheology and sperm velocity were directly comparable. Figure 3 illustrates that sperm velocity decreased as η* increased in a highly correlated manner (p<0.001). A linear model comparing the median log_10_ values of velocity and η* was chosen because the log-transformed data most frequently appeared to be normally distributed according to Anderson-Darling tests. These results imply that the rheology of cervical mucus may be used as surrogate measure of expected sperm penetration.

**Figure 1.**
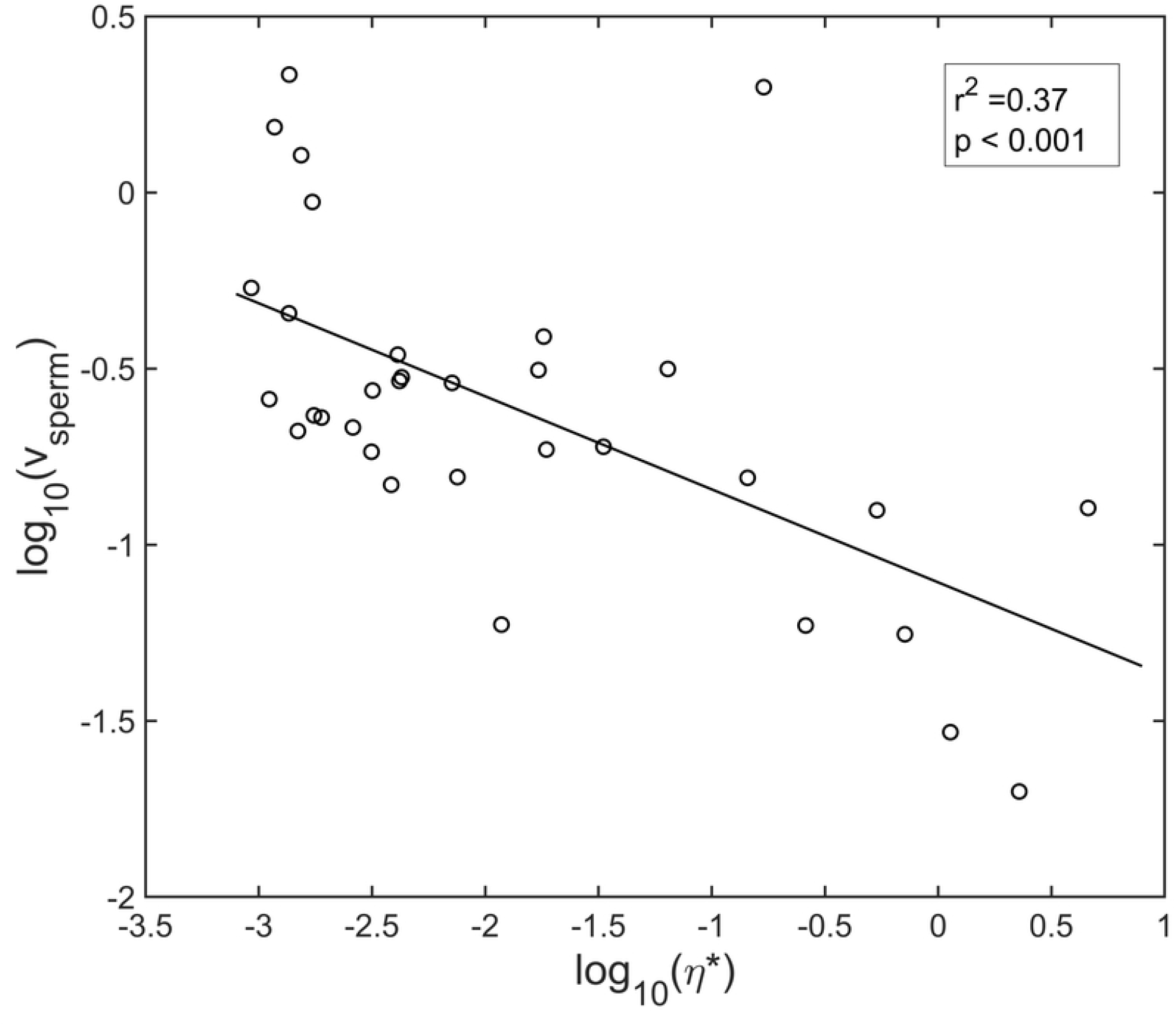
Log_10_ sperm velocity and log_10_ complex viscosity are inversely correlated using linear regression (Pearson, n=6 animals).

## Discussion

Our studies of particle tracking microrheology and sperm motility in the cervical mucus of rhesus macaques have demonstrated that both mucus viscoelasticity and sperm swimming velocity in that mucus are concentration-dependent. Furthermore, we have shown that viscoelasticity as measured by PTMR is correlated with the velocity of sperm in mucus across the range of concentrations studied. This novel finding suggests that PTMR may be used as a surrogate marker of sperm motility in cervical mucus and therefore may be an indicator of the fertility status of the mucus. Additionally, these studies utilize equipment and techniques that could be easily implemented in a clinical research or office environment and better account for the inherent heterogeneity present in both mucus and sperm samples than previously shown. This work represents a step towards more objective and quantitative assessment of fertility attributes of mucus using whole animal samples.

By demonstrating that sperm velocity is reduced by increasing mucus viscosity, our data further supports the existing body of research that designates mucus as a critical regulator of fertility. In the case of existing contraceptives such as progestin-only contraceptives, our findings bolster the notion that a contraceptive effect is likely present through induction of mucus thickening during ovulatory cycles. Our studies may also be useful in understanding fertility changes. Prior to the advent of IUI and IVF, it was thought that up to 15% of women experienced cervical factor infertility, which is infertility related to unfavorable mucus characteristics, even during fertile timepoints. While technologies that physically bypass the cervix have largely rendered this diagnosis moot from a treatment standpoint, there are still clinical scenarios, such as women with cystic fibrosis (CF), where assessment of changes in cervical mucus may help explain the fecundity of an individual. In the specific case of CF, mutations in the CFTR ion channel cause thickened mucus secretions systemically. Today, CF is largely treated with channel modulating drugs.^15^ While the primary focus of these drugs on are the life-extending benefits with correcting CFTR in the lung, other systemic changes, including the cervix, have not been fully characterized and the fertility effect is not totally understood. It is known, however, that pregnancy rates – both planned and unplanned – have increased dramatically in women with CF since the recent advent of modulator therapies,^16^ further implicating cervical mucus hydration as fertility regulator.

Our findings, however, may be most useful for studies designed to augment mucus for the purposes of fertility or contraception. Compared to existing clinical appraisal, PTMR could be employed to provide greater rigor and confidence in assessment of biophysical properties that affect sperm function and, ultimately, sperm ascension. PTMR and sperm tracking serve as probes of these properties in a passive and driven sense, respectively. Measures of sperm penetration represent a logical means to assess fertility-related biophysical properties of cervical mucus. However, existing assays are often limited by poor discrimination power and new technologies such as CASA cannot be used for highly viscous solutions. Meanwhile other rheological evaluations exist. For example, microrheological techniques that evaluate cervical mucus rheology through tracked single particles, sometimes driven by magnets, required much more sophisticated instrumentation.^3^ Additionally, single particle techniques are unable to characterize the dramatic heterogeneity of biophysical properties in individual mucus samples. The innovation represented in this work comes from the combination of techniques that assess and relate sperm and bead motion in cervical mucus via techniques that capture the heterogeneity of both sperm motility and mucus rheology using relatively simple equipment (a microscope and camera).

Our study used macaque whole mucus and sperm samples. These studies need to be validated with human cervical mucus and sperm, though the concentration dependence of mucus from other organ systems like the lung and gut has been consistently demonstrated across non human primate models and in humans, lending confidence to our results in this model.^7^ Additionally, our prior studies demonstrate high amounts of proteomic similarity between macaque and human mucus.^17^ And while there are other mucosal factors (immunological, pH, microbiological, etc.) that affect fertility, from a purely biophysical perspective, recent studies have shown that mucus concentration is the primary regulator of its viscoelasticity.^11,18^ Even so, there may be other factors, including changes in the secreted protein milieu that may affect fertility in ways that we do not account for biophysically.

In conclusion, the cervical mucus barrier regulates fertility as its biophysical properties change in response to hormonal changes throughout the cycle. Quantifying the modulation of those changes due to both endogenous and exogenous perturbation is critical to understanding biological mechanisms and finding therapeutics targeting mucus changes. Non-hormonal compounds are of particular interest for both purposes given the off-target effects that are common with hormone-based drugs. We give an example in closing, knowing that CFTR is hormonally regulated and also implicated in the infertility of women with CF,^16,19^ Figure 4 presents a concept for a prospective contraceptive that inhibits CFTR-mediated anion secretion, resulting in mucus hyperconcentration. Alternatively, a compound that activates ENaC-mediated Na^+^ absorption, may have a similar effect. Recent studies have shown several other ion channels that play a known role in mucus hydration are also present in the cervical epithelium. These may represent drug targets for contraception and infertility, and their ability to induce biophysical changes in the cervical mucosal barrier can be studied via the techniques developed here.

**Figure 4.**
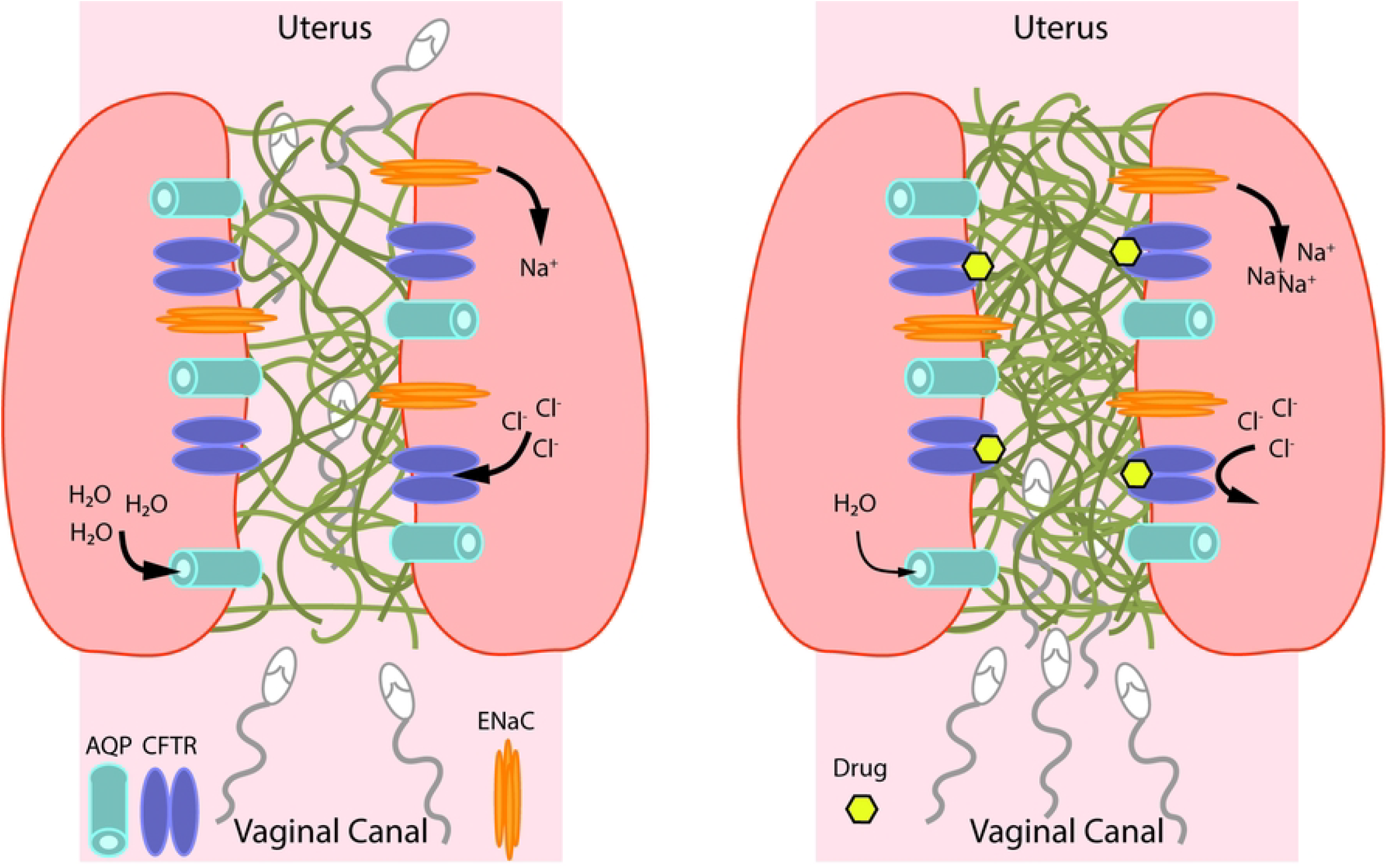
(Left) Liquid and ion transport work in concert to modulate the biophysical properties of fertile peri-ovulatory mucus. The viscoelasticity of the mucin network (green) is sufficiently reduced to allow sperm penetration into the upper reproductive tract. (Right) CFTR inhibition is one possible approach to induce Na+ and liquid absorption to hyperconcentrated and thicken cervical mucus, rendering it impenetrable to sperm.

